# Coro1a promotes the multivesicular body and plasma membrane fusion by facilitating PKM2-mediated SNAP-23 phosphorylation

**DOI:** 10.1101/2025.02.14.638370

**Authors:** Zhenzhai Cai, Zhijie Li, Tingting Yan, Beilei Fu, Xinyu He, Nannan Li, Chenxing Wang, Yujiao Chen, Dingli Zhang, Zhijian Cai, Jing Cai, Weiguo Zhu, Hao Wu

## Abstract

During the formation of endosomal pathway-dependent extracellular vesicles (EVs), intraluminal vesicles in multivesicular bodies (MVBs) fuse with the plasma membrane (PM), releasing intraluminal vesicles to produce exosomes. Soluble N-ethylmaleimide-sensitive factor attachment protein receptor (SNARE) complex, membrane fusion effector, is essential in this process. However, which SNARE complex is involved in MVB and PM fusion and how its assembly is regulated remain elusive. We have demonstrated that neddylation of Coro1a inhibits EV secretion by inducing Rab7-mediated degradation of MVBs. Here, we find that Coro1a increases EV biogenesis by promoting the assembly of SNARE complex STX-12-SNAP-23-VAMP-7 in a neddylation-independent manner. Coro1a activates PKM2 to enhance SNAP-23 phosphorylation in the presence of neddylation inhibitor MLN4924, which drives SNAP-23 to recruit more STX-12 and VAMP7, leading to increased formation of the STX-12-SNAP-23-VAMP-7 complex. Correspondingly, Coro1a-induced EV biogenesis is eliminated by PKM2 inhibitor or SNAP-23 silencing independent of neddylation. Reduced EVs from tumors due to Coro1a knockout cannot be observed in MLN4924-treated tumor mice with PKM2 inhibitor treatment either. Furthermore, Coro1a increases in tumor tissues from lung tumor patients with tumor progression, and high Coro1a indicates unfavorable survival of tumor patients, suggesting that increased Coro1a-mediated enhanced EV production from tumors accelerates tumor progression. Altogether, our data demonstrate that Coro1a facilitates EV biogenesis by promoting the assembly of STX-12-SNAP-23-VAMP-7 complex during MVB and PM fusion independent of neddylation, thereby being involved in the progression of EV-related diseases.

## Introduction

Extracellular vesicles (EVs), roughly classified into ectosomes and exosomes, contain a lot of proteins, nucleic acids, lipids and metabolites from their parental cells, thereby playing an essential role in information communication between different cells ^1^.On the one hand, EVs exert important physiological functions, such as placental and intestinal EVs can mediate fetal immune escape and maintain intestinal immune balance, respectively^2,3^; on the other hand, EVs are involved in the progression of various diseases, including neurodegenerative diseases^4^, cardiovascular diseases^5^ and metabolic diseases^6^. In addition, EVs have pivotal effects on tumor development, especially tumor immunosuppression. Tumor cell-derived EVs (TEVs) profoundly affect the activation of almost all immune effector cells^7^. TEV PD-L1 has been reported to induce systemic immunosuppression^8^. Besides, our previous publication suggests that TEV PD-L1 contributes to anti-PD-L1 therapy resistance by decoying anti-PD-L1^9^. Therefore, targeting TEV production is a prospective strategy to enhance antitumor immunity, which requires further understanding of the mechanisms responsible for EV biogenesis.

Ectosomes originate from outward protrusions of the plasma membrane (PM) that are scissored and shed into extracellular space. Tsg101 is recruited to PM by arrestin domain-containing protein 1 and participates in VPS4-dependent ectosome production. In addition, tetraspanins, including CD9 and CD81, are also responsible for ectosome biogenesis^10^. Exosomes are formed through an endosome-dependent pathway. As the first step of exosome biogenesis, the invagination of PM creates early endosomes. During maturation, the endosomal membrane buds inwardly to form multivesicular bodies (MVBs) rich in intraluminal vesicles (ILVs) ^1^. MVBs mainly have two fates. One is transported to lysosomes for degradation, dominantly driven by Rab7. We have reported that neddylated Coronin-1a (Coro1a) can recruit Rab7 onto MVBs and promote lysosome sorting of MVBs, thereby inhibiting EV biogenesis^11^. The other fate of MVBs is transported to PM for exosome release, which Rab27 dictates predominantly^12^. The soluble N-ethylmaleimide-sensitive fusion attachment protein (SNAP) receptor (SNARE) complexes are responsible for MVB and PM fusion, the last step of exosome biogenesis.

SNARE complexes are vital elements in membrane fusion. SNARE complexes comprise one v-SNARE (vesicle-membrane SNARE) protein and three t-SNARE (target-membrane SNARE) proteins. T-SNARE complexes interact with the v-SNAREs through the N-terminal end of the SNARE motifs, and this nucleates the formation of a four-helical trans-complex, initiating vesicular membrane fusion^13^. SNARE complexes have been demonstrated to regulate EV biogenesis, such as v-SNARE VAMP-3, t-SNARE Syntaxin-3 (STX-3), and SNAP-23, reportedly involved in EV biogenesis^14–16^. However, in addition to the subunits of SNARE complexes, which complete SNARE complex(es) is/are responsible for EV biogenesis and how the assembly of SNARE complexes is regulated during the fusion of MVBs and PM are still largely unknown.

Coro1a is a member of the coronin family associated with F-actin^17^. As mentioned above, we previously reported that Coro1a regulates EV biogenesis by promoting MVB degradation in a neddylation-dependent manner. Furthermore, we found that Coro1a also affects EV biogenesis independent of neddylation^11^. However, the mechanisms of how Coro1a regulates EV biogenesis in a neddylation-independent manner are still undeciphered. Although we investigated the EV biogenesis of the endosomal pathway, the EV we isolated unavoidably mixed with small ectosomes in the study. Therefore, we used the term EVs instead of exosomes to abstain from potential disputes, which was also adopted in this study.

Here, we validate that Coro1a regulates EV biogenesis in a neddylation-independent manner. Overexpression or silencing Coro1a increases or decreases EV release, respectively. Mechanistically, Coro1a promotes PKM2 phosphorylation and enhances the interaction between PKM2 and SNAP-23, leading to increased SNAP-23 phosphorylation. Subsequently, SNAP-23 improves the recruitment of STX-12 and VAMP-7, increases the formation of STX-12-SNAP-23-VAMP-7 complex and strengthens MVB and PM fusion, thereby enhancing EV biogenesis. In addition, we find that Rab27a-mediated regulation of EV biogenesis is mainly Coro1a dependent. Thus, our data reveal a novel mechanism for EV biogenesis involving the regulation of the SNARE complex during MVB and PM fusion.

## Materials and methods

### Mice

Female mice of the C57BL/6J and nude mice, aged between 6-8 weeks, were acquired from Joint Ventures Sipper BK Experimental Animal Co. in Shanghai, China. All the mice were housed in a controlled facility that maintained specific pathogen-free conditions. The experimental procedures on the mice received the necessary approval from the Animal Care and Use Committee of the Zhejiang University School of Medicine.

### Cell lines and cell culture

Human 293T embryonic kidney cells, human HeLa cervical cancer cells and mouse B16F10 melanoma cells were purchased from the Chinese Academy of Sciences Institute (Shanghai, China). The cells were cultured in DMEM with 10% exosome-depleted fetal bovine serum (FBS) (Thermo Fisher Scientific, Waltham, CA, USA) and 1% penicillin/streptomycin (Keyi, Hangzhou, Zhejiang, China). All cells were cultured at 37 °C in a humidified atmosphere with 5% CO_2_.

### Plasmid and DNA, siRNA transfection

Flag-human-*Coro1a*, Myc-human-*Snap23*, His-human-*Pkm2* and SNAP-23 mutants were constructed and acquired from the Miao Ling Plasmid Sharing Platform (Wuhan, Hubei, China). Rab5^Q79L^-GFP was provided by Prof. Yuehai Ke (Zhejiang University, Hangzhou, Zhejiang, China). According to the manufacturer’s protocol, 293T, HeLa and B16F10 cells were transfected with plasmids using JetPEI Transfection Reagent (Polyplus, Beijing, China) or PEI (Polysciences, Warrington, PA, USA) or were transfected with scramble negative control (NC) or target siRNA using INTERFERin (Polyplus). The sequences of the siRNAs used in this study are listed in Supplementary Table 2.

### Semiquantitative EV assay

The method was performed as previously described^11^. Briefly, to detect EVs in the cell culture supernatants, supernatants containing FBS with depleted EVs were cleared by centrifugation at 300 × *g* for 10 min and 2,000 × *g* for 20 min. 4-μm aldehyde sulfate beads (Thermo Fisher Scientific) were first coated with purified anti-CD63 antibodies and then blocked with FBS at room temperature (RT) for 1 h. The beads were washed twice in PBS and centrifuged at 3000 × *g* for 5 min. The cleared supernatants were incubated with anti-CD63-coupled beads overnight at 4°C with shaking. The beads were washed and incubated with anti-CD9 or anti-CD81 antibodies for 30 min at 4°C. After washing, the beads were acquired on an ACEA NovoCyte and the data were analyzed with NovoExpress software (ACEA Biosciences, San Diego, CA, USA). The negative staining threshold was obtained from beads incubated with an unconditioned medium. The antibodies used are listed in Supplementary Table 2.

### Nano-flow cytometry EV assay

EVs isolated from cell culture media, sera or TTs were divided into 50 μl PBS with 1 × 10^10^ particle ml^−1^ concentration. For each 50 μl sample, 0.25 μl specific primary antibodies were added into each sample overnight, and then 0.25 μl secondary antibodies were incubated at 37 °C for 1 h. For FAP staining, EVs were pre-treated with 1 × permeabilization buffer (eBioscience) for 1 h before incubating with a primary antibody. After incubation, the mixture was washed twice with 1 ml PBS by centrifugation at 100,000 g for 20 min at 4 °C. The pellets were resuspended in 100 ml PBS, diluted to the approximate concentration, and analyzed by Flow NanoAnalyzer (N30E, NanoFCM Inc., Xiamen, Fujiang, China).

### Separation of EVs

Cells were seeded onto 15-cm plates at a density of 3 million cells per plate. These cells were cultured for 48 h, and then the media from 10 plates were collected, followed by differential centrifugation. Firstly, the supernatants were centrifuged at 300 × g for 10 min, followed by a centrifugation step at 2,000 × g for 20 min, both carried out at 4°C. Subsequently, another centrifugation was performed at 10,000 × g for 30 min at 4°C. To isolate EVs from tumor tissues (TTs), TTs were detached and ground in 5 ml PBS buffer with a syringe plunger. Then type IV collagenase (Sigma–Aldrich, St. Louis, MO, USA) was added into this buffer at a final concentration of 1 mg ml^−1^ and enzymatically digested for 1 h at 37 °C. The TT fragments containing supernatants were centrifuged at 300 × g for 10 min. Then, all the resulting supernatants were filtered through 0.22 μm syringe filters (Millipore, Billerica, MA, USA) and collected in 35 ml ultracentrifuge tubes (Beckman Coulter, Brea, CA, USA). EVs were concentrated by ultracentrifugation using an SW32Ti rotor (L-90K with SW32Ti rotor, Beckman Coulter) at 100,000 × g for 70 min at 4°C. Subsequently, the EV pellets were resuspended in sterile PBS. The protein content of the isolated EVs was quantified using BCA protein assay without adding any detergent (Thermo Fisher Scientific).

### Transmission electron microscopy (TEM) and nanoparticle tracking analysis (NTA)

EV suspension was placed on 200-mesh carbon-coated copper grids at RT for 2 min. Any excess suspension was carefully removed from the grids using filter paper. Subsequently, the EVs were subjected to negative staining by uranyl acetate, performed at RT for 5 min. After the staining step, the EVs were washed twice with PBS and left to dry. The dried EVs were then observed using an FEI Tecnai T10 electron microscope operating at 100 kV (Thermo FEI, Hillsboro, OR, USA).

To measure particle sizes and concentrations, NTA analyzed EVs using a NanoSight NS300 system (Malvern PANalytical, Malvern, UK) configured with a 488 nm laser and high-sensitivity sCMOS camera.

### Immunoprecipitation and western blotting

EVs and cells were lysed in the SDS buffer. The lysates were then boiled at 100 °C for 10 min. Following the lysis and boiling step, the samples were resolved using SDS-polyacrylamide gel and transferred onto PVDF membranes (Millipore). These membranes were blocked with 5% milk for 2 h. Next, the membranes were incubated overnight with the appropriate primary antibodies at 4 °C, followed by incubation with the corresponding secondary antibodies at RT for 2 h. An Enhanced Chemiluminescence Kit (MultiSciences, Hangzhou, Zhejiang, China) was employed to detect the protein bands.

For immunoprecipitation (IP), the cells were lysed in co-immunoprecipitation lysis buffer consisting of 50 mM Tris-HCl, 5 mM EDTA, 150 mM NaCl, 0.5% (v/v) NP-40, and 10% (v/v) glycerol at pH 7.4. The lysis buffer was supplemented with 1 mM PMSF, 1 mM Na_3_VO_4_, and 10 mM NaF. The lysate was then incubated with M2-Flag beads (Sigma–Aldrich) overnight at 4 °C for IP. The immunoprecipitates were washed at least three times with lysis buffer and subsequently analyzed using the indicated antibodies by western blotting. The Abs used are listed in Supplementary Table 2.

### Iodixanol density gradient fractionation

Iodixanol density gradient fractionation was performed as previously described^11^. Iodixanol density gradient medium (StemCell, Vancouver, BC, Canada) was prepared in ice-cold PBS immediately before use to generate a discontinuous step gradient (12%-28%).The 28% iodixanol solution was added to the bottom of a centrifuge tube, followed by carefully layered iodixanol solutions of decreasing concentrations in PBS to form the gradient. Identical gradients without samples were prepared and ultracentrifuged to determine fraction densities, measured by refractometry. Crude EV pellets resuspended in ice-cold PBS were loaded on top of the gradient. The gradient was subjected to ultracentrifugation at 120,000 × *g* for 16 h at 4°C using Beckman XPN-100 with an SW 41 Ti swinging bucket rotor. Nine individual 1-ml fractions were collected from the top of the gradient. For western blotting analysis, each individual 1 ml fraction was transferred to a new ultracentrifugation tube, diluted in PBS and subjected to ultracentrifugation at 120,000 × *g* for 4 h at 4 °C using an SW 32 Ti swinging bucket rotor. The resulting pellets were lysed in 5 × SDS buffer on ice and boiled for 10 min at 100 °C.

### Immunofluorescence

The cells were fixed with precooled methyl alcohol for 10 min at −20 °C and then permeabilized by 0.1% Triton X-100 for 10 min at RT. Non-specific binding sites were blocked by incubating the cells with a blocking buffer containing 5% bovine serum albumin and 3% goat serum in PBS. Then, the cells were incubated with the corresponding primary antibodies overnight at 4 °C in the blocking buffer. After 3 washes in PBS the following day, the cells were incubated with DyLight 488-labeled secondary antibodies for 1 h at RT and washed in PBS. Finally, nuclei were stained with DAPI (Thermo Fisher Scientific). The stained sections were imaged using an Olympus IX83-FV3000 confocal microscope (Olympus Corp, Tokyo, Japan). Images were analyzed with ImageJ software (NIH, Bethesda, MD, USA). The antibodies used are listed in Supplementary Table 2.

### CRISPR-Cas9 mediated construction of gene knockout cells

The gRNAs, with sequences for their target genes listed in Supplementary Table 2, were annealed and cloned into the Lenti-CRISPR-v2 vector. To delete target genes, HeLa and B16F10 were transiently transfected with Lenti-CRISPR-v2 plasmid carrying target gRNAs and selected with 2.5 µg/ml puromycin for 3 days. Cells were then transferred into fresh medium without puromycin and seeded at super-low density to allow colony formation from single cells. Colonies were then picked and expanded for knockout validation by immunoblot.

### Total internal reflection fluorescence (TIRF) microscopy

TIRF microscopy was performed as previously described^11^. Briefly, HeLa cells were transfected with the GFP-CD63-His or mCherry-CD63-HA plasmid for 36 h and incubated in PBS before analysis by TIRF microscopy. For TIRF microscopy with an Olympus IX83 microscope (Olympus), the penetration depth (δ) of the evanescent field used to excite the fluorophore was set to 150 nm. Frames were acquired at 10 Hz in stacks of 400 images with an exposure time of 100 ms. Fluorescence image acquisitions were collected with CellSens software (Olympus). GFP^+^ or mCherry^+^ CD63 vesicles were quantified using ImageJ software (NIH). To analyze vesicle motion, time-series images were captured with a Nikon N-STORM & A1 Cell TIRF system using a DU897 EMCCD 100× oil TIRF objective and the fluorescence image acquisitions were collected with Nis-Elements software (Nikon, Tokyo, Japan). Imaris 9.5 software (Bitplane AG, Zurich, Switzerland) calculated the mean square displacement. The diffusion coefficient, the slope of the linear fit of the first 15 points of the mean square displacement curve, was then calculated as *D_xy_*= s/4.

### Lipid-mixing assay

The lipid-mixing assay was performed according to previous publication with minor modifications^18^. Return the lipid powder stored at −20 °C to room temperature. POPC/DOPS was dissolved with chloroform, and DiD/DiI was dissolved with ethanol. Make T-liposome (2.5 μM total lipids, POPC:DOPS:DiD = 78%:20%:2%) and V-liposome (2.5 μM total lipids, POPC:DOPS:DiI = 78%:20%:2%). The lipid mixture is blown dry in nitrogen, which involves tilting the glass tube until a thin film forms on the wall. The glass tube is then placed in a light-avoiding vacuum desiccator to dry thoroughly. Add 400 μl Buffer A (50 mM Tris-Cl pH 7.5, 150 mM NaCl, 0.5 mM TCEP, 50 mM beta-OG) to the dried lipid films and the vortex mixture then clears the solution. Mix 5 nM purified STX-12 full-length proteins with a lipid-detergent mixture containing POPC, DOPS, and DiD. Transfer the mixtures into a tube and incubate in the dark at room temperature for 20 min. Mix 5 nM purified VAMP7Δ12-3M proteins with a lipid-detergent mixture containing POPC, DOPS, and DiI, and then transfer the mixtures into a tube and incubate in the dark at room temperature for 20 min. The final volume of the protein-lipid-detergent mixture should be less than the maximum capacity of the desalting column. Load the protein-lipid-detergent mixtures into the Sephadex G-25 desalting column(GE Healthcare,America). The colored fractions were collected. 100 μM t-liposome harboring STX-12 and 100 μM v-liposome harboring VAMP7 are mixed with the addition of 5 μM SNAP-23 to a final volume of 70 μl with Buffer B (50 mM Tris-Cl pH 7.5, 150 mM NaCl,0.5 mM TCEP) in the well of a standard 96-well plate. Liposome fusion analysis was performed using the FluoDia T70 fluorescent flat screen Reader (Protein Technologies, America) equipped with a 530/10 excitation filter, 580/10 and 667/10 emission filter at 37 °C. Donor (DiI) and acceptor (DiD) fluorescence were monitored every 25 seconds. Monitoring the fluorescence of the donor liposome (DiI) and recipient liposome (DiD) can calculate the original FRET efficiency, proximity ratio, in DiD and DiI, indicating the liposome fusion signal: EPR = IDiD/(IDiD + IDiI).

### Protein purification

The recombinant proteins in this study were expressed in E. coli BL21 DE3. In general, cells were cultured to optical density 600 nm (OD 600 nm) = 0.4-0.8 and induced by 0.1 mM isopropyl-β-d-thiogalactoside at 100rpm for 18-24 h at 18 °C. 1.2 l of cultured cells were collected by centrifugation at 7,000 g for 5 min. Cell pellets were resuspended by lysis buffer (1M Tris-Cl pH7.5, 2.5M NaCl) supplied with 1M dithiothreitol, 1 mM PMSF and 1% tritonx-100. Resuspended cells were broken by sonication at 40 W on ice for 10 min and centrifuged at 9,500 rpm for 30 min at 4 °C to separate supernatants and cell debris. 1 ml of Ni-NTA prepacked column(Sangon Biotech,China) washed by PBS and imidazole for 10 column volumes. Then, the collected supernatant was rotated in the column and bonded overnight at 4 °C.On the second day, the PBS was first rotated and combined in the column for 10 min at 4 °C, then washed with 10-column volume PBS. Then, imidazole with different concentration gradients was washed out of the column, and effluent samples were collected.

### Mass spectrum (MS)

To analyze the proteins interacting with SNAP-23, ectopically expressed SNAP-23 was pulled down from 293T cell lysates by anti-Myc-coated beads were separated by SDS-PAGE and stained with Coomassie brilliant blue R-250 (Solarbio, Beijing, China). Then, the gel was cut into 1 cm × 0.5 cm pieces. The protein content was analyzed by MS using the Q Exactive system.

### Tumor model and treatment

On day 0, mice were subcutaneously injected with 2 × 10^6^ tumor cells (B16F10, B16F10 *Coro1a^−/−^* cells) or 1 × 10^7^ tumor cells (HeLa and HeLa *Coro1a^−/−^* cells). On day 7, the tumor-bearing mice were randomized into groups and received treatments as described below. The B16F10 and B16F10 *Coro1a^−/−^* tumor-bearing mice were intratumorally injected with DMSO or Shikonin (2 mg/kg/injection/2 days) for 4 injections. The tumor mice also received an intratumoral injection with MLN4924 (2 mg/kg/injectin/2 days) for 4 injections. Tumor growth was monitored every 2 days by measuring the length and width of the tumors. Tumor volume was calculated using the formula: 0.5 × length × width^2^.

### Statistical analysis

Data are expressed as the mean ± s.d.. The normal distribution of data was validated by the Shapro-Wilk test. Unpaired Student’s *t*-test was used to compare two groups and one-way ANOVA for comparisons among multiple groups using Graph Prism 8.0 software (GraphPad Software Inc., San Diego, CA, USA). Statistical significance was considered if a *P* value was < 0.05.

## Results

### Coro1a enhances EV release by promoting PM and MVB fusion

We previously reported that Coro1a regulates EV biogenesis in both neddylation-dependent and neddylation-independent manners. However, we did not reveal the mechanisms responsible for the neddylation-independent regulation of EV biogenesis by Coro1a^11^. To elucidate this issue, we overexpressed Coro1a in HeLa cervical cancer cells in the presence of MLN4924, a specific inhibitor of E1 NEDD8-activating enzyme that blocks neddylation modification. Corresponding to our previous findings, we found that ectopic expression of Coro1a still increased the release of CD9^+^CD63^+^ and CD81^+^ CD63^+^ EVs from MLN4924-treated 293T and HeLa cells (Fig. 1A). This effect of Coro1a was further confirmed by the increases in protein content and particle number of the EVs (Supplementary Fig. 1A, B). Correspondingly, Coro1a overexpression increased the signals for several EV-enriched markers, including CD63, Alix, Tsg101 and CD81, in EVs from the same number of MLN4924-treated 293T cells with or without iodixanol gradient purification (Fig. 1B and Supplementary Fig. 1C). However, Coro1a affected neither the viability of MLN4924-treated 293T and HeLa cells nor the size distribution and morphology of EVs from 293T cells (Supplementary Fig. 1D-F).

**Figure 1.**
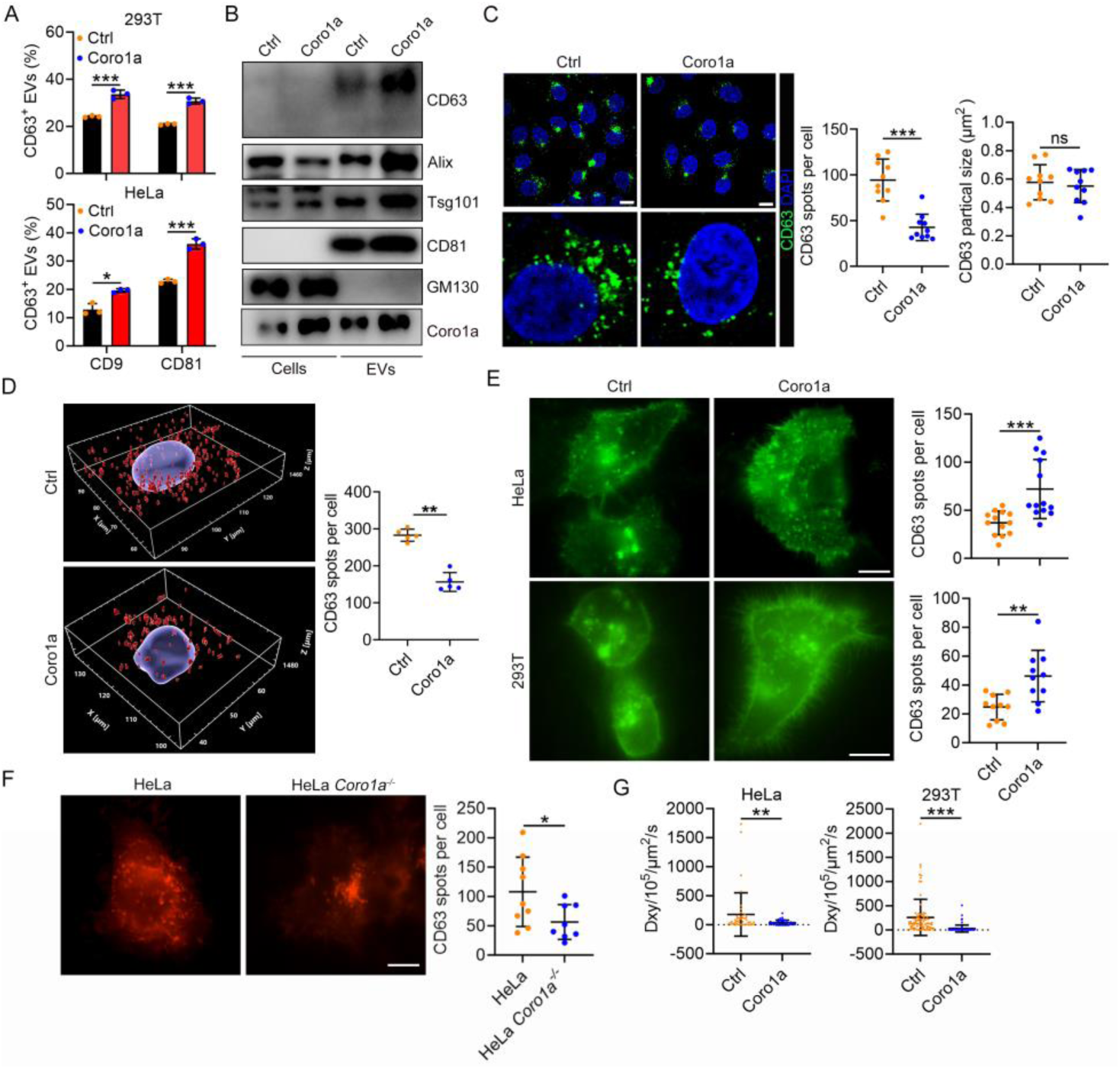
Coro1a enhances EV release by promoting PM and MVB fusion. (A) 293T and HeLa cells with or without Coro1a overexpression were treated with 100 nM MLN4924 for 12 h. EVs in the supernatants were captured with anti-CD63-coated beads, and then CD9^+^CD63^+^ or CD81^+^CD63^+^ EVs were analyzed by flow cytometry (n = 3). (B) EVs were isolated from equal numbers of the corresponding 293T cells in A. Western blotting analysis to detect the indicated EV markers. (C) Confocal microscopy analysis of the MVB marker CD63 in the corresponding HeLa cells in A; CD63 spot numbers and particle size were statistically analyzed (n = 10). Scale bar, 10 µm. (D) 3D quantification of CD63^+^ MVBs in the corresponding HeLa cells in A by z-stack confocal microscope; CD63 spot numbers and particle size were statistically analyzed (n = 5). (E-G) TIRF microscopy analysis of the CD63^+^ MVB distribution in cells in A (E) or HeLa and HeLa *Coro1a^−/−^*cells in the presence of MLN4924 (F); CD63 spot numbers in the subplasmalemmal region were statistically analyzed (n = 13). Scale bar, 10 μm. (G) Mean diffusion coefficient (*Dxy*) values for the individual trajectories of MVBs in F are shown (n = 42 in HeLa and n = 83 in 293T). Representative results from three independent experiments are shown. ns, not significant; \**P* < 0.05; \*\**P* < 0.01 and \*\*\**P* < 0.001 (unpaired two-tailed Student’s *t*-test; mean ± s.d.; except for unpaired Mann-Whitney test and median ± interquartile ranges in F).

Then, we sought to uncover which step(s) of EV biogenesis is(are) affected by Coro1a in a neddylation-independent manner. We first investigated the effects of Coro1a on CD63^+^ MVBs. Coro1a-overexpressing decreased CD63^+^ MVBs in MLN4924-treated HeLa cells but had minimal effect on their size (Fig. 1C). Similar results were obtained in HRS^+^ MVBs (Supplementary Fig. 1G). Moreover, 3D quantification of CD63^+^ MVBs in z-stack confocal images further confirmed this phenotype (Fig. 1D). Subsequently, we examined the impact of Coro1a on MVB origin and degradation. Coro1a did not alter the numbers of EEA1^+^ early endosomes (EEs) in MLN4924-treated HeLa cells (Supplementary Fig. 1H), the first step of MVB biogenesis. In addition, when EEs were enlarged by overexpression of constitutively active Rab5^Q79L^, we observed a minimal effect of Coro1a on the budding of CD63^+^ intraluminal vesicles (Supplementary Fig. 1I). These results indicate that non-neddylated Coro1a does not affect the MVB formation. We also found that Coro1a did not affect the colocalization of CD63^+^ MVBs and LAMP1^+^ lysosomes in MLN4924-treated HeLa cells (Supplementary Fig. 1J), suggesting the unchanged degradation of MVBs.

Since Coro1a showed no effects on EV biogenesis and degradation, we assumed whether Coro1a increases EV secretion by promoting MVB and PM fusion. If so, Coro1a should induce MVB docking onto the PM. Next, we used total internal reflection fluorescent (TIRF) microscopy to evaluate PM docking of MVBs. To monitor the CD63^+^ MVBs, we transfected HeLa cells with plasmids expressing CD63 fluorescent fusion protein and then transfected control or Coro1a into these cells. In contrast to the reduced CD63^+^ MVBs in total cells, Coro1a increased CD63^+^ MVBs in the subplasmalemmal region of MLN4924-treated HeLa and 293T cells (Fig. 1E). Then, we transfected plasmids expressing CD63 fluorescent fusion protein into HeLa cells without and with Coro1a knockout (HeLa *Coro1a^−/−^* cells), and confirmed the comparable transfection efficacy of CD63 (Supplementary Fig. 1K). We found the opposite result between MLN4924-treated HeLa cells with or without Coro1a knockout (Fig. 1F and Supplementary Fig. 1L). As the first step of vesicle fusion, docking is crucial^19^. After PM docking, the motion of vesicles is restricted^20^. To confirm whether non-neddylated Coro1a enhances EV production by increasing the docking of MVBs and PM, the coefficient *D_xy_* of CD63^+^ MVBs, an index of mobility, was calculated along vesicle trajectories using a rolling analysis window. Both *D_xy_* in MLN4924-treated HeLa and 293T cells significantly declined after Coro1a overexpression (Fig. 1G). Altogether, these results demonstrate that Coro1a promotes the fusion of MVBs and PM, thereby increasing EV release in a neddylation-independent manner.

### Coro1a induces the formation of STX-12-SNAP-23-VAMP-7 complex

SNARE complexes are essential to regulate exocytosis by promoting vesicle fusion with PM^21^. SNARE proteins, including v-SNARE VAMP-3 and t-SNAREs STX-3 and SNAP-23^14–16^, have been demonstrated to regulate EV secretion. Therefore, we individually silenced VAMP-3, STX-3 and SNAP-23 and found that SNAP-23 but not VAMP-3 or STX-3 silencing abolished Coro1a-mediated increase in CD81^+^CD63^+^ EV secretion from MLN4924-treated cells (Fig. 2A and Supplementary Fig. 2A, B). SNAP-23 silencing also reduced CD9^+^CD63^+^ EV secretion from MLN4924-treated 293T cells (Fig. 2A). Similar results were also obtained in MLN4924-treated HeLa cells (Fig. 2B). In addition, decreased EV secretion from MLN4924-treated HeLa *Coro1a^−/−^* cells was no longer detected after SNAP-23 silencing (Supplementary Fig. 2C). Then, we found that ectopic expression of Coro1a remarkably induced SNAP-23 distribution in CD63^+^ MVBs of MLN4924-treated HeLa cells (Fig. 2C). Therefore, non-neddylated Coro1a regulates EV biogenesis via SNAP-23.

**FIGURE 2.**
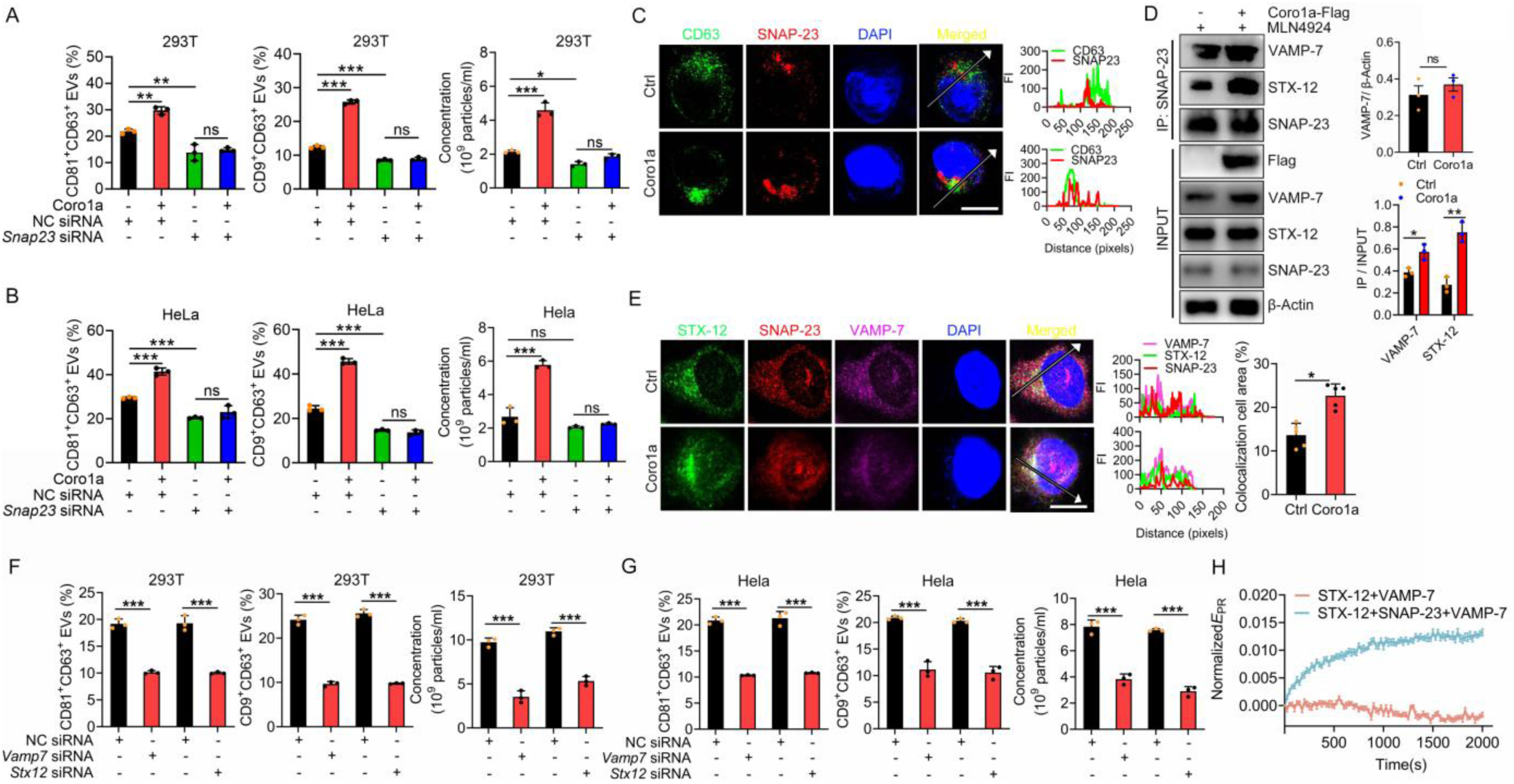
Coro1a induces the formation of the STX-12-SNAP-23-VAMP-7 complex. (A, B) 293T (A) or HeLa (B) cells with or without Coro1a overexpression were transfected with NC or *Snap23* siRNA for 24 h. Then, the cells were treated with 100 nM MLN4924 for 12 h, and the EVs in the supernatants were captured with anti-CD63-coated beads, followed by flow cytometric analysis of CD9^+^CD63^+^ or CD81^+^CD63^+^ EVs (n = 3), and nano-flow cytometry was employed to measure the EV concentration (n = 3). (C) Confocal microscopy analysis of the colocalization of CD63 and SNAP-23 in MLN-treated HeLa cells with or without Coro1a overexpression. (D) Western blotting analysis of STX-12 and VAMP-7 in the lysates of MLN-treated 293T cells with or without Coro1a overexpression after IP with anti-SNAP-23. (E) Confocal microscopy analysis of STX-12, VAMP-7 and SNAP-23 colocalized in MLN-treated HeLa cells with or without Coro1a overexpression (left), and the colocalizations of STX-12, VAMP-7 and SNAP-23 were statistically analyzed (right, n = 5). Representative results from three independent experiments are shown. Scale bar, 10 μm. (F, G) 293T (F) or HeLa (G) cells were transfected with NC, *Vamp7* or *Stx12* siRNA for 48 h. Then, the EVs in the supernatants were captured with anti-CD63-coated beads, followed by flow cytometric analysis of CD9^+^CD63^+^ or CD81^+^CD63^+^ EVs (n = 3), and nano-flow cytometry was employed to measure the EV concentration (n = 3). (H) Time-dependent liposome fusion was measured from the development of FRET between the DiD-labeled STX-12 liposomes and the DiI-labeled VAMP-7Δ12-3 M liposomes. Each trace is the average from three independent replicates. ns, not significant; \*\**P* < 0.01 and \*\*\**P* < 0.001 (unpaired two-tailed Student’s *t*-tes; mean ± s.d.).

To elucidate the SNARE complex responsible for MVB and PM fusion, we immunoprecipitated ectopic expressed SNAP-23 in MLN4924-treated 293T cell lysates, performed MS analysis and found that SNARE proteins, including STX-12 and VAMP-7, were pulled down by SNAP-23 (Supplementary Fig. 2D). We found that Coro1a overexpression caused more VAMP-7 and STX-12 immunoprecipitated by SNAP-23 in MLN4924-treated 293T cells (Fig. 2D). The opposite results were obtained in MLN4924-treated HeLa *Coro1a^−/−^* cells (Supplementary Fig. 2E). The Coro1a-induced increase in the colocalization of SNAP-23, VAMP-7 and STX-12 in MLN4924-treated HeLa cells was also confirmed by immunofluorescence staining (Fig. 2E). Furthermore, VAMP-7 or STX-12 silencing significantly reduced CD81^+^CD63^+^ and CD9^+^CD63^+^ EV secretion by 293T and HeLa cells (Fig. 2F, G and Supplementary Fig. 2F). We next conducted lipid-mixing experiments to explore whether VAMP-7, STX-12 and SNAP-23 form a functional SNARE complex to drive MVB-PM fusion. Since VAMP-7 favors a closed conformation with its N-terminal longin domain bound to its SNARE motif, we introduced Y45E/I139S/I144S mutation (3 M) to facilitate the opening of VAMP-7^22–24^ and eliminated the C-terminal 12-amino acids to improve VAMP-7 solubility. With lipid-mixing assays, we observed significant membrane fusion between VAMP-7Δ12-3 M liposomes and STX-12 liposomes in the presence of SNAP-23 (Fig. 2H), demonstrating that the SNARE complex composed of STX-12, SNAP-23 and VAMP-7, is competent in mediating MVB-PM fusion. These results suggest that Coro1a promotes STX-12-SNAP-23-VAMP-7 complex formation in a neddylation-independent manner.

### Coro1a facilitates PKM2-mediated SNAP-23 phosphorylation

Next, we wondered how non-neddylated Coro1a regulates the formation of the STX-12-SNAP-23-VAMP-7 complex. PKM2 is involved in EV biogenesis by inducing Serine/Threonine phosphorylation (p-Ser/Thr) of SNAP-23^16^. MS results revealed that PKM2 was also pulled down by SNAP-23 (Supplementary Fig. 2D). We did detect the interaction between PKM2 and SNAP-23 in 293T cells, which was not affected by MLN4924 (Fig. 3A). So we explored the role of PKM2 in non-neddylated Coro1a-mediated regulation of the formation of the STX-12-SNAP-23-VAMP-7 complex. First, we found that Coro1a induced SNAP-23 phosphorylation in MLN4924-treated 293T cells (Fig. 3B). On the contrary, lowered phosphorylated SNAP-23 was detected in MLN4924-treated HeLa *Coro1a^−/−^* cells (Supplementary Fig. 3A). Consistent with its function on EV biogenesis, we found that constitutive-active SNAP-23 mutant (SNAP-23-S95E) remarkably promoted CD81^+^CD63^+^ EV release by 293T cells, but dominant-negative SNAP-23 mutant (SNAP-23-S95A) did not show this effect (Fig. 3C). In contrast, SNAP-23-S95A started to inhibit CD81^+^CD63^+^ EV release with higher overexpression (Supplementary Fig. 3B). Furthermore, we found that SNAP-23-S95E overexpression abolished the difference in CD81^+^CD63^+^ EV secretion between MLN4924-treated HeLa *Coro1a^−/−^* and HeLa cells (Fig. 3D). These results indicate that Coro1a regulates neddylation-independent EV secretion by affecting p-Ser/Thr of SNAP-23. Then, we found that PKM2 inhibitor Shikonin eliminated Coro1a-induced SNAP-23 phosphorylation (Fig. 3E). Correspondingly, Shikonin commensurated the total numbers of CD63^+^ MVBs between MLN4924-treated HeLa *Coro1a^−/−^* and HeLa cells (Fig. 3F, G). Furthermore, we found comparable STX-12-SNAP-23-VAMP-7 complexes in these cells (Fig. 3H). Similar results were obtained by PKM2 silencing (Supplementary Fig. 3C-E). MLN4924 has been reported to increase glycolysis by activating PKM2 via promoting its tetramerization^25^. To exclude the interference of MLN4924 on this process, we inhibited neddylation by NEDD8 silencing and found that PKM2 silencing abolished Coro1a-induced p-Ser/Thr of SNAP-23 in 293T cells with NEDD8 silencing (Supplementary Fig. 3F). Furthermore, Coro1a deficiency-induced decrease in SNAP-23 interaction with VAMP-7 and STX-12 was also negated in NEDD8-silenced HeLa cells along with PKM2 silencing (Supplementary Fig. 3G). Subsequently, we found a similar subplasmalemmal distribution of CD63^+^ MVBs between Shikonin-treated HeLa *Coro1a^−/−^* and HeLa cells in the presence of MLN4924 (Fig. 3I). Therefore, Coro1a increases SNAP-23 phosphorylation by facilitating PKM2 activation.

**FIGURE 3.**
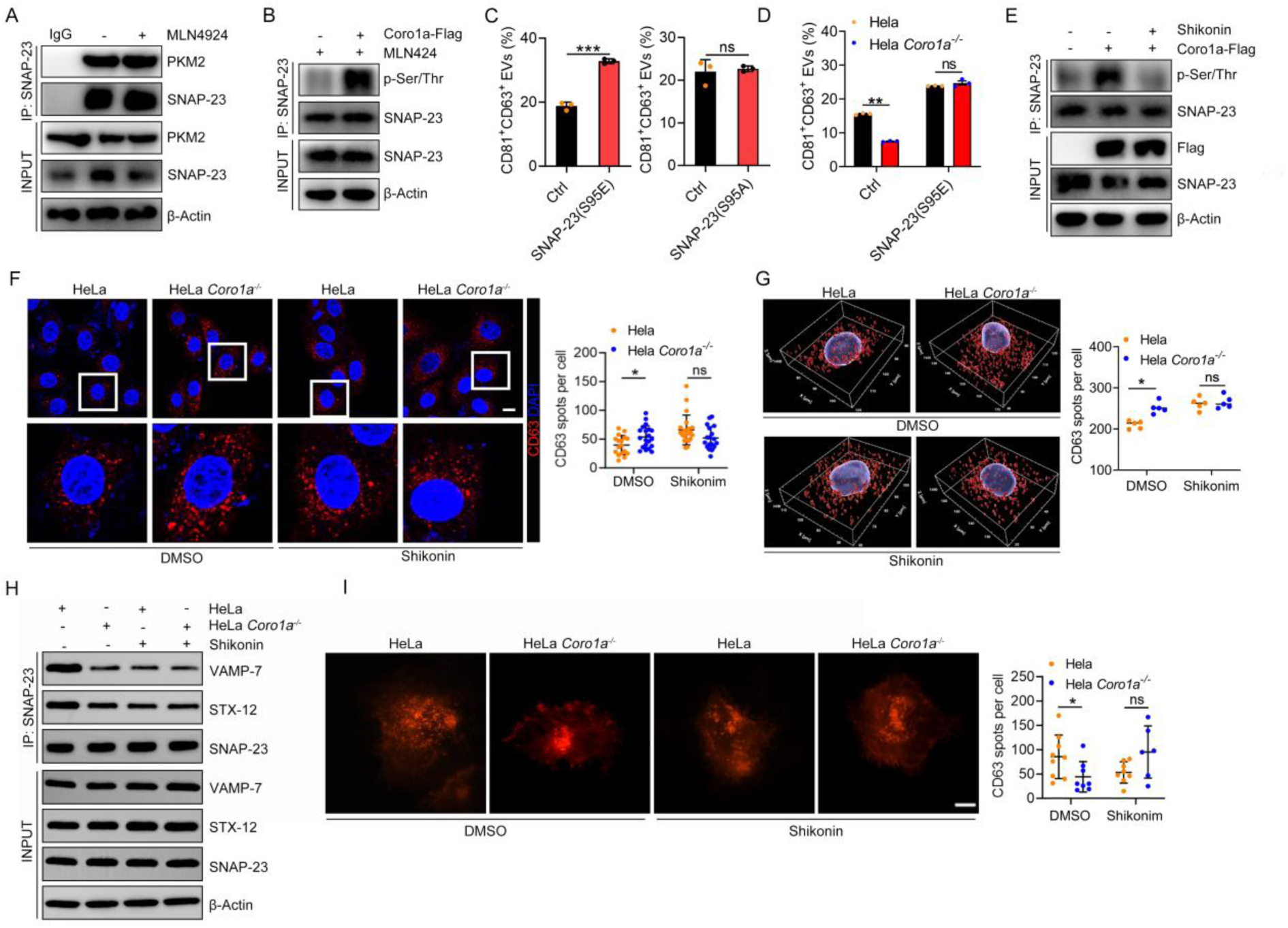
Coro1a facilitates PKM2-mediated SNAP-23 phosphorylation. (A) Western blotting analysis of PKM2 in the lysates of 293T cells with or without 100 nM MLN4924 for 12 h after IP with anti-SNAP-23. (B) Western blotting analysis of p-Ser/Thr of SNAP-23 in the lysates of MLN4924-treated 293T cells with or without Coro1a overexpression after IP with anti-SNAP-23. (C) 293T cells were transfected with SNAP-23-S95E or SNAP-23-S95A for 48 h. Then, the EVs in the supernatants were captured with anti-CD63-coated beads, followed by flow cytometric analysis of CD81^+^CD63^+^ EVs (n = 3). (D) MLN4924-treated HeLa *Coro1a^−/−^* and HeLa cells were transfected with SNAP-23-S95E for 48 h. Then, the EVs in the supernatants were captured with anti-CD63-coated beads, followed by flow cytometric analysis of CD81^+^CD63^+^ EVs (n = 3). (E) Western blotting analysis of p-Ser/Thr of SNAP-23 in the lysates of MLN4924-treated 293T cells with or without Coro1a overexpression, accompanied by 1 μM Shikonin treatment for 12 h after IP with anti-SNAP-23. (F) Confocal microscopy analysis of CD63 in Shikonin-treated HeLa and HeLa *Coro1a^−/−^* cells in the presence of MLN4924; CD63 spot numbers were statistically analyzed (n = 20). (G) 3D quantification of CD63^+^ MVBs in Shikonin-treated HeLa and HeLa *Coro1a^−/−^* cells in the presence of MLN4924 by z-stack confocal microscope; CD63 spot numbers and particle size were statistically analyzed (n = 5). (H) Western blotting analysis of VAMP-7 and STX-12 in Shikonin-treated HeLa and HeLa *Coro1a^−/−^* cells in the presence of MLN4924 after IP with anti-SNAP-23. (I) TIRF microscopy analysis of the CD63^+^ MVB distribution in Shikonin-treated HeLa and HeLa *Coro1a^−/−^* cells in the presence of MLN4924; CD63 spot numbers in the subplasmalemmal region were statistically analyzed (n = 6-10). Scale bar, 10 μm. Representative results from three independent experiments are shown. ns, not significant; \**P* < 0.05; \*\*\**P* < 0.001 (unpaired two-tailed Student’s *t*-test; mean ± s.d.).

### PKM2 dictates Coro1a-induced EV biogenesis

Subsequently, we investigated the functional relevance of PKM2 to non-neddylated Coro1a-mediated regulation of EV biogenesis. PKM2 inhibitor or silencing eliminated Coro1a overexpression-mediated increase in CD81^+^CD63^+^ EV release from MLN4924-treated HeLa cells (Fig. 4A, B). Coro1a overexpression also could not increase CD81^+^CD63^+^ EV release from NEDD8 knockout HeLa (HeLa *Nedd8^−/−^*) cells in the presence of PKM2 inhibitor (Supplementary Fig. 4A, B). A similar result was observed in MLN4924-treated 293T cells along with Shikonin treatment (Supplementary Fig. 4C). On the contrary, Shikonin negated Coro1a knockout-mediated decrease in CD81^+^CD63^+^ EV release from MLN4924-treated HeLa cells (Supplementary Fig. 4D). Then, we examined the role of non-neddylated Coro1a in regulating EV production *in vivo*, we established murine subcutaneous HeLa tumors and B16F10 melanoma with or without Coro1a knockout (Supplementary Fig. 4E) and simultaneously treated these mice with MLN4924. We found that EV amounts in TTs from both Coro1a knockout tumor-bearing mice were significantly reduced relative to mice inoculated with wild-type tumors. However, these differences were abolished in both tumor-bearing mice receiving Shikonin treatment (Fig. 4C, D). Shikonin also abrogated the decrease in TT EV amount of B16F10 *Coro1a^−/−^*; *Nedd8^−/−^*tumor-bearing mice relative to B16F10 *Nedd8^−/−^* tumor-bearing mice (Supplementary Fig. 4F, G). Previous and our publications have demonstrated that robust antitumor immunity can be induced by reducing tumor EV production, thereby inhibiting tumor growth^8,11^. Consistent with the above results, the B16F10 tumors exhibited obvious growth priority relative to B16F10 *Coro1a^−/−^* tumors in immune complete mice, while both tumors showed similar development in Shikonin-treated mice (Fig. 4E). Similar results were obtained in B16F10 *Nedd8^−/−^* and B16F10 *Coro1a^−/−^*; *Nedd8^−/−^*tumor-bearing mice with Shikonin treatment (Supplementary Fig. 4H). In addition, when EVs from B16F10 cells (B16F10-EVs) were supplied to B16F10 *Coro1a^−/−^* tumor-bearing mice, comparable tumor growth was observed between MLN4924-treated B16F10 and B16F10 *Coro1a^−/−^* tumor-bearing mice (Fig. 4F), suggests that reduced TEV production led to Coro1a deficiency-mediated tumor suppression. Thus, these results suggest that Coro1a regulates EV biogenesis by affecting PKM2 activation.

**FIGURE 4.**
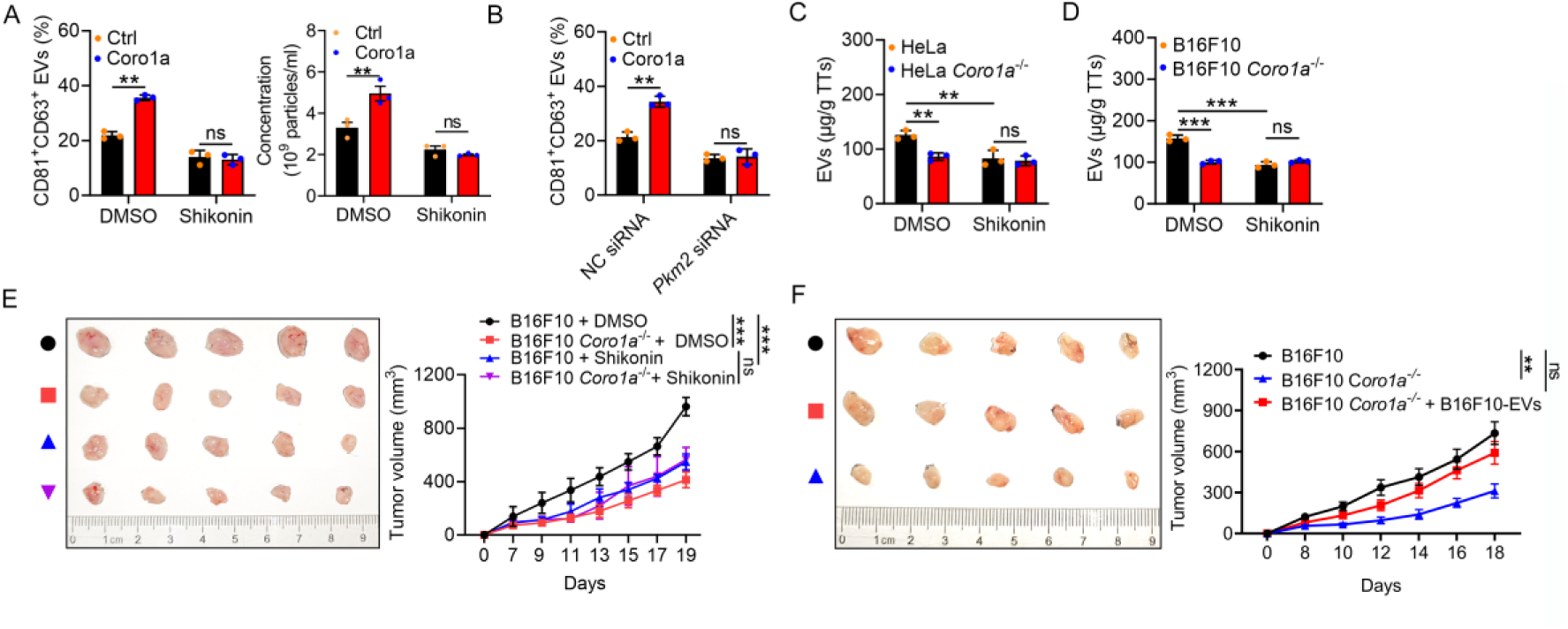
PKM2 dictates Coro1a-induced EV biogenesis. (A, B) HeLa cells with or without Coro1a overexpression were treated with Shikonin (A) or transfected with *Pkm2* siRNA (B). Then, the cells were treated with 100 nM MLN4924 for 12 h, and the EVs in the supernatants were captured with anti-CD63-coated beads, followed by flow cytometric analysis of CD81^+^CD63^+^ EVs (n = 3), and nano-flow cytometry (A) was employed to measure the EV concentration (n = 3). (C, D) Nude mice were inoculated with HeLa and HeLa *Coro1a^−/−^* cells for 3 weeks (C) or B16F10 and B16F10 *Coro1a^−/−^* cells for 2 weeks (D). TTs were collected, and EV amounts were quantified (n = 3). (E) After being inoculated with B16F10 and B16F10 *Coro1a^−/−^* cells for 7 days, the tumor-bearing mice were intratumorally injected with DMSO or Shikonin (2 mg/kg/injection/2 days). The tumor mice also received an intratumoral injection with MLN4924 (10 mg/kg/injectin/2 days). The tumor size was monitored every other day (n = 5). (F) After being inoculated with B16F10 and B16F10 *Coro1a^−/−^* cells for 7 days, the tumor-bearing mice received an intratumoral injection with MLN4924 (10 mg/kg/injectin/2 days). Some B16F10 *Coro1a^−/−^* tumor-bearing mice were intravenously supplied with 20 μg of B16F10-EVs 3 times per week for 2 weeks. The tumor size was monitored every other day (n = 5). Representative results from two independent experiments are shown. ns, not significant; \*\**P* < 0.01 and \*\*\**P* < 0.001 (Unpaired two-tailed Student’s *t*-test in A, B; One-way ANOVA followed by Turkey test in C-E; mean ± s.d.).

### Coro1a strengthens the interaction between PKM2 and SNAP-23

Next, we wondered how Coro1a regulates PKM2-mediated SNAP-23 phosphorylation. First, we found that Coro1a interacted with PKM2 rather than SNAP23 in 293T cells independently of neddylation (Fig. 5A). PKM2 is particularly likely to form a dimer when tyrosine 105 (Y105) is phosphorylated^26^. Then, dimerized PKM2 is re-located at the vesicular structures close to the PMs, associating with SNAP-23^16^. We found that Coro1a induced PKM2 phosphorylation at the Y105 site in the absence of Coro1a neddylation (Fig. 5B). Consistently with this result, Coro1a increased exogenous and endogenous interaction between PKM2 and SNAP-23 (Fig. 5C, D). In contrast, PKM2 and SNAP-23 interaction was inhibited in HeLa *Coro1a^−/−^* cells (Fig. 5E). Therefore, Coro1a increases SNAP-23 and PKM2 interaction by promoting PKM2 phosphorylation.

**FIGURE 5.**
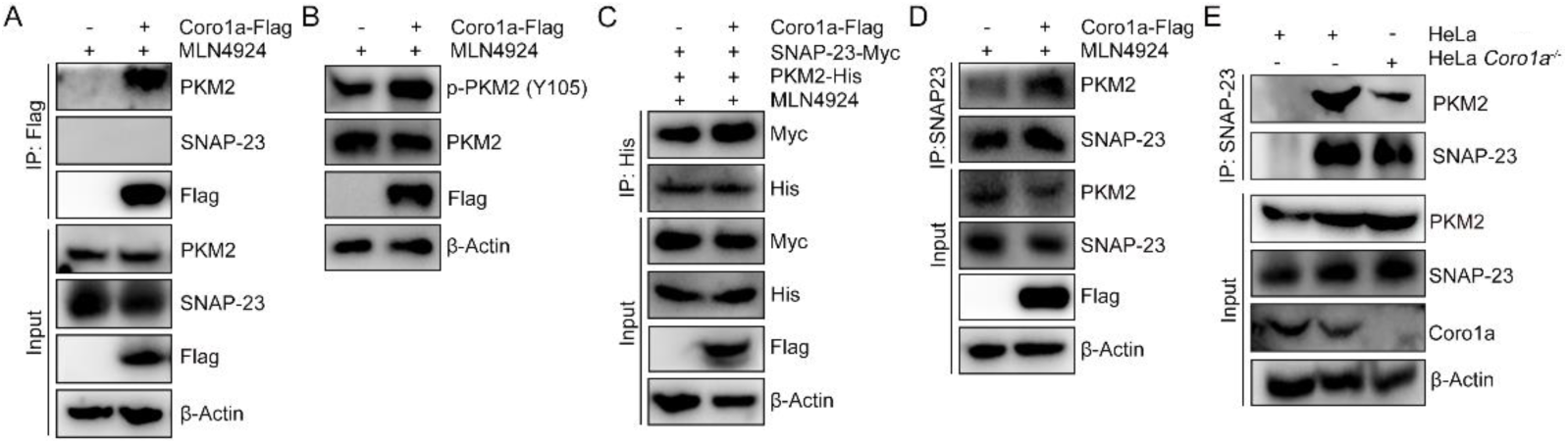
Coro1a strengthens the interaction between PKM2 and SNAP-23. (A) Western blotting analysis of PKM2 and SNAP-23 in the lysates of MLN4924-treated 293T cells with or without Coro1a-Flag overexpression after IP with anti-Flag. (B) Western blotting analysis of Y105 phosphorylation of PKM2 in MLN4924-treated 293T cells with or without Coro1a overexpression. (C) Western blotting analysis of Myc in the lysates of MLN4924-treated 293T cells with or without Coro1a-Flag overexpression, accompanied by PKM2-His and SNAP-23-Myc overexpression after IP with anti-His. (D) Western blotting analysis of PKM2 in the MLN4924-treated 293T cell lysates with or without Coro1a-Flag overexpression after IP with anti-SNAP-23. (E) Western blotting analysis of PKM2 in the lysates of MLN4924-treated HeLa and HeLa *Coro1a^−/−^* cells after IP with anti-SNAP-23. Representative results from two independent experiments are shown.

### Advanced tumors increase EV release by upregulating Coro1a

Tumors secrete more EVs as progression^27^. We found sequentially increased Coro1a in TTs of 2-, 3- and 4-week tumor mice bearing B16F10 tumors and increased EVs from these TTs (Fig. 6A and Supplementary Fig. 5A). The increase in Coro1a of these TTs was further confirmed by immunohistochemistry (IHC) (Fig. 6B). In addition, increased extent of EVs in TTs were greatly blunted with Coro1a deficiency (Fig. 6C), suggesting tumors secret more EVs as progression by upregulating Coro1a.

**Figure 6.**
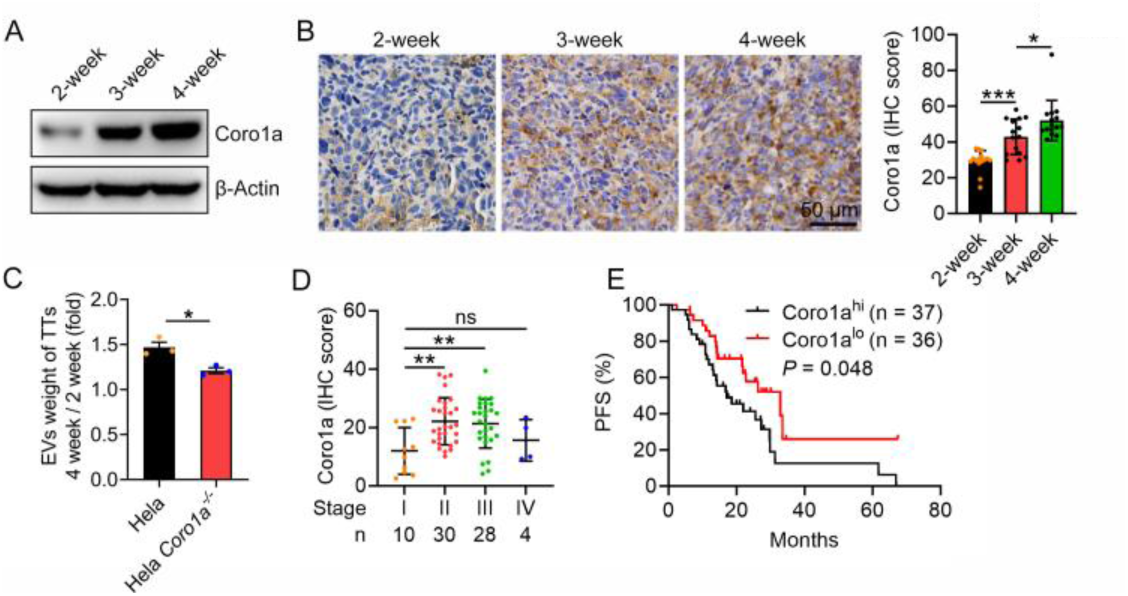
Advanced tumors increase EV release by upregulating Coro1a. (A, B) Coro1a proteins in TTs from mice bearing B16F10 tumors for 2, 3 and 4 weeks were detected by western blotting (A) and IHC (B). (C) Nude mice were inoculated with HeLa and HeLa *Coro1a^−/−^* cells for 2 or 4 weeks. TTs were collected, and EV amounts were quantified (n = 3). (D, E) Coro1a proteins in TTs from lung cancer patients were detected by IHC. Coro1a protein levels in patients at different stages were statistically analyzed (D), and the PFS of Coro1a^hi^ and Coro1a^lo^ patients (E). Scale bar, 50 μm. ns, not significant; \**P* < 0.05 (unpaired two-tailed Student’s *t*-test in C; Log-rank test in D; mean ± s.d.).

As an extension of our findings in human cancers, we observed significantly higher Coro1a in cancerous (Ca) tissues of non-small cell lung cancer (NSCLC) patients than in precancerous (Para-Ca) tissues (Supplementary Fig. 5B and Supplementary Table 1). Compared with Stage I patients, Coro1a protein levels in TTs of Stage II and III patients were significantly high. However, this increase was not obtained in Stage IV patients, probably due to the small sample size (Fig. 6D). We previously demonstrated that Coro1a deficiency inhibited mouse MC38 colon tumor growth and stimulated antitumor CD8^+^ T cell responses, which were abolished by supplementation of TEVs, indicating Coro1a accelerates tumor growth by suppressing antitumor immunity through inducing TEV production^11^. Consistent with the oncogenesis role of TEVs and the regulatory effect of Coro1a on EVs, we found that patients with low Coro1a protein levels (Coro1a^lo^) had a favorable progress-free survival (PFS) relative to patients with high Coro1a protein levels (Coro1a^hi^) (Fig. 6E). These results suggest that increased TEVs are produced in a Coro1a-dependent manner, which aggravates tumor progression.

## Discussion

Accumulating evidence supports the notion that EVs play crucial roles in the progression of multiple diseases. Therefore, pathogenic EVs are fascinating intervention targets for treating the corresponding diseases. However, current strategies to reduce EV production are still unsuitable for clinical application. To achieve this purpose, further exploring the mechanisms for EV biogenesis is necessary, which will benefit the finding of specific promising targets for intervention. Previously, we found that neddylated Coro1a is located on the MVB membrane and can recruit Rab7 onto MVBs by direct binding to Rab7. Subsequently, Rab7 guides the MVBs towards a fate of lysosomal degradation, leading to the inhibition of EV biogenesis. However, we also found that in the presence of MLN4924, Coro1a knockout still suppressed EV secretion, suggesting a neddylation-independent regulation of EV biogenesis by Coro1a^11^. Here, we revealed that non-neddylated Coro1a increases EV secretion by promoting the fusion of MVB and PM. Mechanistically, Coro1a activates PKM2, enhances PKM2-mediated SNAP-23 phosphorylation, and promotes STX-12-SNAP-23-VAMP-7 complex assembly and subsequent MVB and PM fusion, thereby facilitating EV release. So, Coro1a can regulate EV biogenesis in both neddylation-dependent and neddylation-independent manners. However, we previously demonstrated that in the absence of MLN4924, Coro1a still increased EV production^11^, which indicates that non-neddylated Coro1a-mediated regulation in EV biogenesis is dominant and EV biogenesis inhibited by neddylated Coro1a is likely strengthened in responses to certain external stresses. Regardless, Coro1a vividly represents the requirement for dedicated regulation of EV biogenesis.

Coro1a promotes PKM2-mediated SNAP-23 phosphorylation. However, we detected Coro1a bound PKM2 rather than SNAP-23, suggesting that Coro1a increases SNAP-23 phosphorylation not by strengthening the interaction between PKM2 and SNAP-23. PKM2 has dimer and tetramer forms. Dimerized PKM2 possesses low pyruvate kinase activity but high protein serine/threonine kinase activity^28^. Phosphorylation of PKM2 at the Y105 site inhibits tetramerization of PKM2, thereby increasing its protein kinase activity^29^. We found that Coro1a promoted the phosphorylation of PKM2 at the Y105 site, which is consistent with the result that increased serine/threonine phosphorylation of SNAP-23 is induced by Coro1a. Coro1a contains no kinase domain, and as Ser/Thr kinase, PKM2 cannot induce auto-tyrosine phosphorylation. We assumed that Coro1a likely facilitates the interaction of a tyrosine kinase with PKM2, leading to increased PKM2 phosphorylation. The tyrosine kinase involved in this process needs further study.

We previously demonstrated that Coro1a deficiency greatly delayed the growth of murine MC38 colon tumors by inhibiting EV-mediated antitumor immunity^11^. In this study, we found that Coro1a increased in tumors as tumor progression. Consistent with the function of Coro1a on EV secretion, we detected enhanced EV amount from advanced tumors. In addition, we found that patients with high Coro1a protein levels had an unfavorable PFS. TEVs are quite immunosuppressive and responsible for the malignancy of tumors^8,30^. Therefore, increased Coro1a with tumor progression probably shortened the PFS of tumor patients by promoting the production of TEVs. However, we did not identify the factor(s) mediating Coro1a upregulation in tumors as progression. It is worth elucidating this issue, which will benefit the finding of a new target for tumor therapy.

In summary, we elucidate how Coro1a regulates EV biogenesis through a neddylation-independent pathway in this study, extending our understanding of the dual effects of Coro1a on EV biogenesis. In addition, we also reveal a novel mechanism for EV biogenesis, which may contribute to developing potential strategies to reduce pathogenic EVs in the future.

## Acknowledgments

This work was supported by the Plan Project of Wenzhou Municipal Science and Technology (Y20210883), Academician Helin’s new Medical Research Fund (19331104) and the Project of Zhejiang Education Department (202045547).

## Author contributions

Z.Z.C., Z.J.L., T.T.Y., B.L.F. X.Y.H., N.N.L., C.X.W., Y.J.C., and D.L.Z. performed various experiments; Z.J.C. discussed the manuscript. J.C., W.G.Z. and H.Wu. designed the project and supervised the study; Z.Z.C. and Z.J.L. wrote the manuscript.

## Data availability statement

The data for the study are available from the corresponding author upon reasonable request.

